# Tension-dependent regulation of mammalian Hippo signaling through LIMD1

**DOI:** 10.1101/238394

**Authors:** Consuelo Ibar, Elmira Kirichenko, Benjamin Keepers, Edward Enners, Katelyn Fleisch, Kenneth D. Irvine

**Affiliations:** Waksman Institute and Department of Molecular Biology and Biochemistry, Rutgers University, Piscataway NJ 08854 USA.

## Abstract

Hippo signaling is regulated by biochemical and biomechanical cues that influence the cytoskeleton, but the mechanisms that mediate this have remained unclear. We show that all three mammalian Ajuba family proteins – AJUBA, LIMD1, and WTIP - exhibit tension-dependent localization to adherens junctions, and that both Lats family proteins, LATS1 and LATS2, exhibit an overlapping tension-dependent junctional localization. This localization of Ajuba and Lats family proteins is also influenced by cell density, and by Rho activation. We establish that junctional localization of Lats kinases requires LIMD1, and that LIMD1 is also specifically required for the regulation of Lats kinases and YAP by Rho. Our results identify a biomechanical pathway that contributes to regulation of mammalian Hippo signaling, establish that this occurs through tension-dependent LIMD1-mediated recruitment and inhibition of Lats kinases in junctional complexes, and identify roles for this pathway in both Rho-mediated and density-dependent regulation of Hippo signaling.

## INTRODUCTION

The Hippo signaling network integrates a wide range of biochemical and biomechanical cues to modulate organ growth and cell fate (Irvine and Harvey, 2015; Meng et al., 2016). This control of growth and cell fate is effected through regulation of two homologous transcriptional co-activator proteins, YAP and TAZ. Many of the upstream cues that influence YAP and TAZ activity converge on cell junctions and the actin cytoskeleton (Dupont, 2016; Sun and Irvine, 2016). For example, YAP and TAZ can be affected by cell-cell contacts (Kim et al., 2011), cell-extracellular matrix contacts (Kim and Gumbiner, 2015; Serrano et al., 2013; Zhao et al., 2012), cell density (Zhao et al., 2007), F-actin levels (Aragona et al., 2013; Fernández et al., 2011; Sansores-Garcia et al., 2011), cytoskeletal tension (Dupont et al., 2011; Rauskolb et al., 2014), cell shape (Dupont et al., 2011; Wada et al., 2011), substrate stiffness (Dupont et al., 2011), cell stretching (Aragona et al., 2013; Benham-Pyle et al., 2015; Codelia et al., 2014) and Rho activity (Dupont et al., 2011; Yu et al., 2012). However, we have only a very limited understanding of the molecular processes that mediate this responsiveness to the cytoskeleton and mechanical stresses.

A major, though not exclusive, mechanism of YAP and TAZ regulation is through phosphorylation by the Lats kinases, LATS1 and LATS2 (Meng et al., 2016; Sun and Irvine, 2016). Phosphorylation of YAP and TAZ by Lats kinases inhibits their activity by promoting their degradation, and their exclusion from the nucleus. Lats kinases are activated through phosphorylation by upstream kinases, including Mst and Map4k family kinases. Hippo signaling was first identified in *Drosophila* based on the overgrowth phenotypes associated with mutations in *warts* and upstream pathway components that promote activation of Warts, which is the *Drosophila* orthologue of the Lats kinases (Reddy and Irvine, 2008). These overgrowths occur in response to abnormally elevated levels of Yorkie activity (the orthologue of YAP and TAZ) (Huang et al., 2005). Similarly, elevated YAP and TAZ activity is observed in many cancers (Harvey et al., 2013).

Studies in *Drosophila* identified a mechanism for biomechanical regulation of Hippo signaling involving tension-dependent recruitment of Warts into a complex at adherens junctions with the *Drosophila* Ajuba family protein Jub (Rauskolb et al., 2014). Jub, which contributes to regulation of Hippo signaling during development and regeneration (Das Thakur et al., 2010; Meserve and Duronio, 2015; Sun and Irvine, 2011), is recruited to adherens junctions in a tension-dependent manner (Rauskolb et al., 2014). Jub is an inhibitor of Warts (Das Thakur et al., 2010), and recruitment of Warts to Jub complexes also prevents it from localizing to other junctional and apical complexes where Warts activation occurs (Su et al., 2017; Sun et al., 2015). Whether a comparable mechanism exists in mammalian cells has been disputed (Jagannathan et al., 2016), and studies of biomechanical regulation of Hippo signaling have focused on other potential cues, including tension at focal adhesions, actin levels and organization, and mechanically-gated channels (Dupont, 2016).

Mammals have three Ajuba family proteins: AJUBA, WTIP, and LIMD1. They have been ascribed a variety of cellular localizations, including cytoplasm, nucleus, centrosomes, adherens junctions, focal adhesions, and P bodies (Goyal et al., 1999; Hirota et al., 2003; Kanungo et al., 2000; Kim et al., 2012; Marie et al., 2003; Pratt et al., 2005; Spendlove et al., 2008; Srichai et al., 2004). They have also been ascribed a wide variety of biological functions, but one key function identified for Ajuba family proteins is physical interaction with, and inhibition of, Lats kinases (Abe et al., 2006; Das Thakur et al., 2010). The association of Ajuba family proteins with Lats kinases can be enhanced by JNK or ERK phosphorylation (Reddy and Irvine, 2013; Sun and Irvine, 2013). However, aside from one report implicating LIMD1 in a JNK-dependent activation of YAP after cyclic stretch (Codelia et al., 2014), it has not been shown that mammalian Ajuba family proteins contribute to biomechanical regulation of Hippo signaling. Indeed, a recent report has suggested that Ajuba proteins do not participate in biomechanical regulation of Hippo signaling, and that they interact with Lats kinases exclusively in the cytoplasm (Jagannathan et al., 2016). We also note that while association of LIMD1 with focal adhesions is reduced by blebbistatin treatment (Schiller et al., 2011), indicating that it depends upon myosin activity, whether LIMD1 localization to adherens junctions is also tension-dependent has not been investigated, nor has any contribution of cytoskeletal tension to AJUBA, WTIP, LATS1 or LATS2 localization been reported.

Here, we describe investigations of the biomechanical regulation of Ajuba family proteins, and their contribution to Hippo signaling. We find that AJUBA, WTIP, and LIMD1 each exhibit a strong tension-dependent association to adherens junctions. We show that both LATS1 and LATS2 (henceforth collectively referred to as LATS) also exhibit a tension-dependent localization to adherens junctions. In MCF10A cells, one of the three Ajuba family proteins, LIMD1, is specifically required for the junctional localization of LATS. Using pharmacological inhibition of cytoskeletal tension, cell density, and Rho activation as models of cytoskeletal regulation of Hippo signaling, we show that LIMD1 is specifically required for cytoskeletal regulation of YAP, and that this regulation correlates with recruitment of LATS into complexes at adherens junctions. Our results indicate that LIMD1 is essential to certain modes of biomechanical regulation of Hippo signaling, and functions by recruiting LATS into an inhibitory complex at adherens junctions.

## RESULTS

### Tension-dependent localization of Ajuba family proteins in mammalian epithelial cells

To investigate regulation of Ajuba family protein localization in epithelial cells, we created constructs expressing GFP-tagged human cDNAs in a lentiviral vector with a doxycycline (Dox)-inducible promoter (CMV-TetON). We then used these to express Ajuba family proteins in the canine kidney epithelial cell line MDCKIIG, and in the human breast epithelial cell line MCF10A. Cells with relatively homogeneous expression of each construct were selected after viral transduction. Using this system, proteins can be expressed at low levels by using low doses of Dox (Table 1), thereby minimizing or eliminating mis-localization due to over-expression. Western blotting cell lysates confirmed inducible expression for each protein, and in cases where we had antibodies that recognize endogenously expressed proteins on Western blots (AJUBA and LIMD1), was used to identify conditions under which expression of GFP-tagged proteins is comparable to endogenous expression levels (Fig. S1d, e).

**Table 1.**
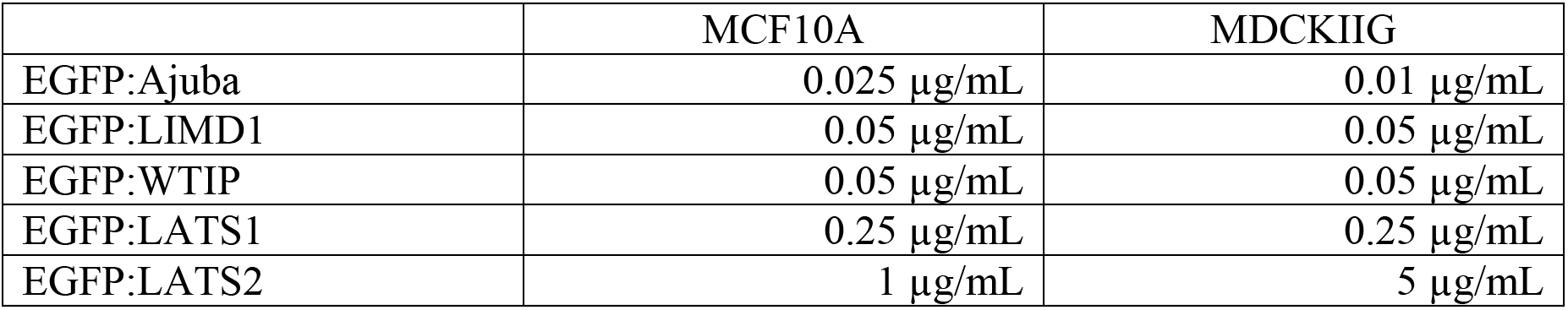
Amounts of doxycycline used to induce the different cell lines for confocal images.

We found that in low density cultures with cells in contact, each of the three Ajuba family proteins exhibit a prominent, punctate localization to adherens junctions (identified by overlap with the apical circumferential localization of E-cadherin (E-cad)) (Figs. 1, S2a-d) and a faint localization to the cytoplasm. Some localization to focal adhesions, identified by overlap with basal puncta of Vinculin (VCL), could also be detected, but at lower levels than at adherens junctions, and this focal adhesion localization was more evident in MDCKIIG cells than in MCF10A cells (Fig. S3). In MDCKIIG cells, which have linear adherens junctions, Ajuba family proteins preferentially accumulate in puncta near intercellular vertices (Figs. 1, S2b, d, f). This contrasts with the more even distribution of E-cad or ZO-1 around the cell circumference (Fig. 1b, d, S2b, d), and is reminiscent of the distribution of Jub in *Drosophila* imaginal disc epithelia (Rauskolb et al., 2014). In MCF10A cells, which have punctate adherens junctions (Meng and Takeichi, 2009), Ajuba family proteins have a punctate distribution all around the apical cell periphery that matches the distribution of the junctional markers E-cad and ZO-1 (Figs. 1a,b, S2a,c). We also attempted to examine the localization of endogenous Ajuba family proteins using commercially available antisera. Antisera against AJUBA and LIMD1 that detect junctional staining in MCF10A and MDCKIIG cells similar to that of GFP-tagged proteins were identified (Fig. S1).

**Figure 1.**
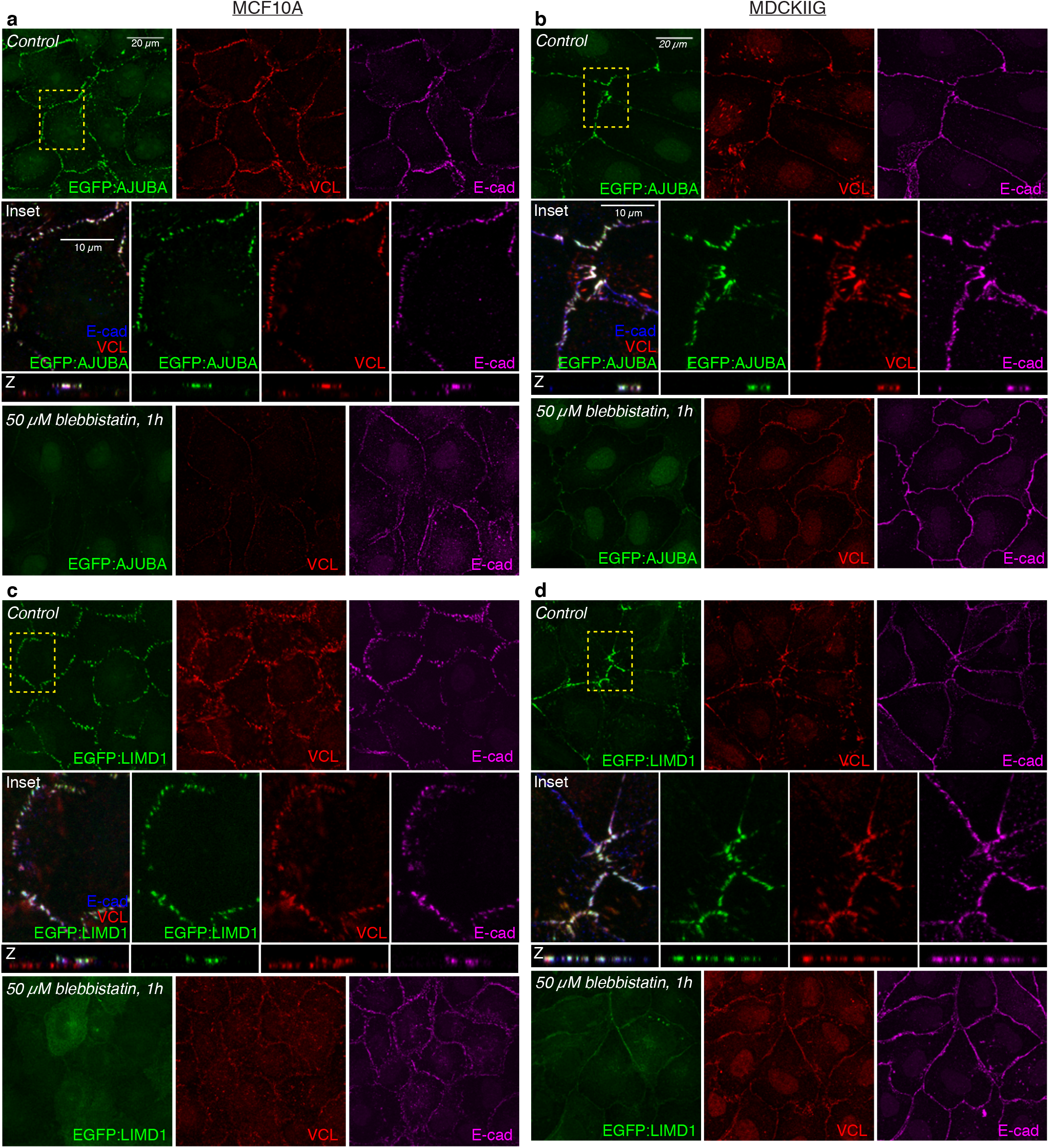
Tension-dependent localization of EGFP:Ajuba family proteins in stable MCF10A or MDCKIIG cell lines. MCF10A **(a, c)** or MDCKIIG **(b, d)** cells from the indicated cell lines were plated at low-density (15,000 cells/cm^2^) and cultured for 48 hours in total. Transgene expression was induced with Dox (Table 1) for 24 hours before treatments. Cells were treated with DMSO (Control) or 50 μM blebbistatin for 1 hour, fixed in the presence of 0.5% Triton X-100 and then stained with VCL (red) and E-cadherin (E-cad, magenta). Square images are apical slices for MCF10A and Z-projections for MDCKIIG cells, and are representative of at least 3 biological replicates. Insets show higher magnification of the boxed regions (yellow dashes). Vertical confocal slices (Z) are shown at the same magnification as the insets.

In addition to its localization to focal adhesions, VCL exhibits a tension-dependent localization to adherens junctions (le Duc et al., 2010; Yonemura et al., 2010). Notably, VCL localization overlaps with Ajuba protein family localization at adherens junctions, including both the localization to punctate adherens junctions in MCF10A cells, and the localization to dispersed puncta in MDCKIIG cells (Figs. 1, S2b). This co-localization suggests that like VCL, Ajuba family localization to adherens junctions is dependent upon cytoskeletal tension. To confirm this, we treated cells either with either of two inhibitors of myosin activity: the Rho-associated protein kinase (ROCK) inhibitor Y-27632, or the Myosin II inhibitor blebbistatin. These compounds substantially reduce cytoskeletal tension, and treatment of cells with either compound led to a loss of the bright puncta of AJUBA, LIMD1, or WTIP, both at adherens junctions and at focal adhesions, and in both MDCKIIG and MCF10A cells (Figs 1, S2). In MDCKIIG cells, but not in MCF10A cells, we also detected a low level of Ajuba family protein distribution along cell junctions that is evenly distributed around the cell perimeter after inhibiting cytoskeletal tension (Figs 1b,d, S2b,d,f). Thus each of the three Ajuba family proteins has a localization profile that is modulated by cytoskeletal tension, with strong punctate junctional localization, and also focal adhesion localization, dependent upon cytoskeletal tension.

α-Catenin has been implicated in tension-dependent recruitment of VCL and Jub to adherens junctions, which is thought to occur through a tension-dependent change in α-Catenin conformation (Yonemura et al., 2010). α-Catenin has also been reported to bind to AJUBA, but the potential influence of tension on this interaction was not investigated (Marie et al., 2003). To assess the requirement for α-Catenin in tension-dependent recruitment of mammalian Ajuba family proteins, we examined LIMD1 localization in cells with an siRNA-mediated reduction in α-Catenin expression. This substantially reduced junctional levels of LIMD1 (Fig. 2a, b). To further analyze potential interaction between LIMD1 and α-Catenin, we used proximity ligation assays (PLA) (Fredriksson et al., 2002) to probe their physical proximity in vivo. A robust and specific PLA signal was detected using antibodies against α-Catenin and GFP in EGFP:LIMD1-expressing cells, and this signal was abolished by blebbistatin treatment (Fig. 2d). As PLA is thought to require proteins to be within 30-40 nm, this signal indicates that LIMD1 and α-Catenin are closely associated on cell-cell junctions. The tension-dependent conformational change in α-Catenin exposes the epitope recognized by the a18 monoclonal antibody (Yonemura et al., 2010). Junctional puncta of LIMD1, AJUBA, and WTIP co-localize with a18 staining in both MCF10A and MDCKIIG cells (Fig. 2c). Together, these observations suggest that LIMD1 is recruited to adherens junctions by the tensed, open conformation of α-Catenin.

**Figure 2.**
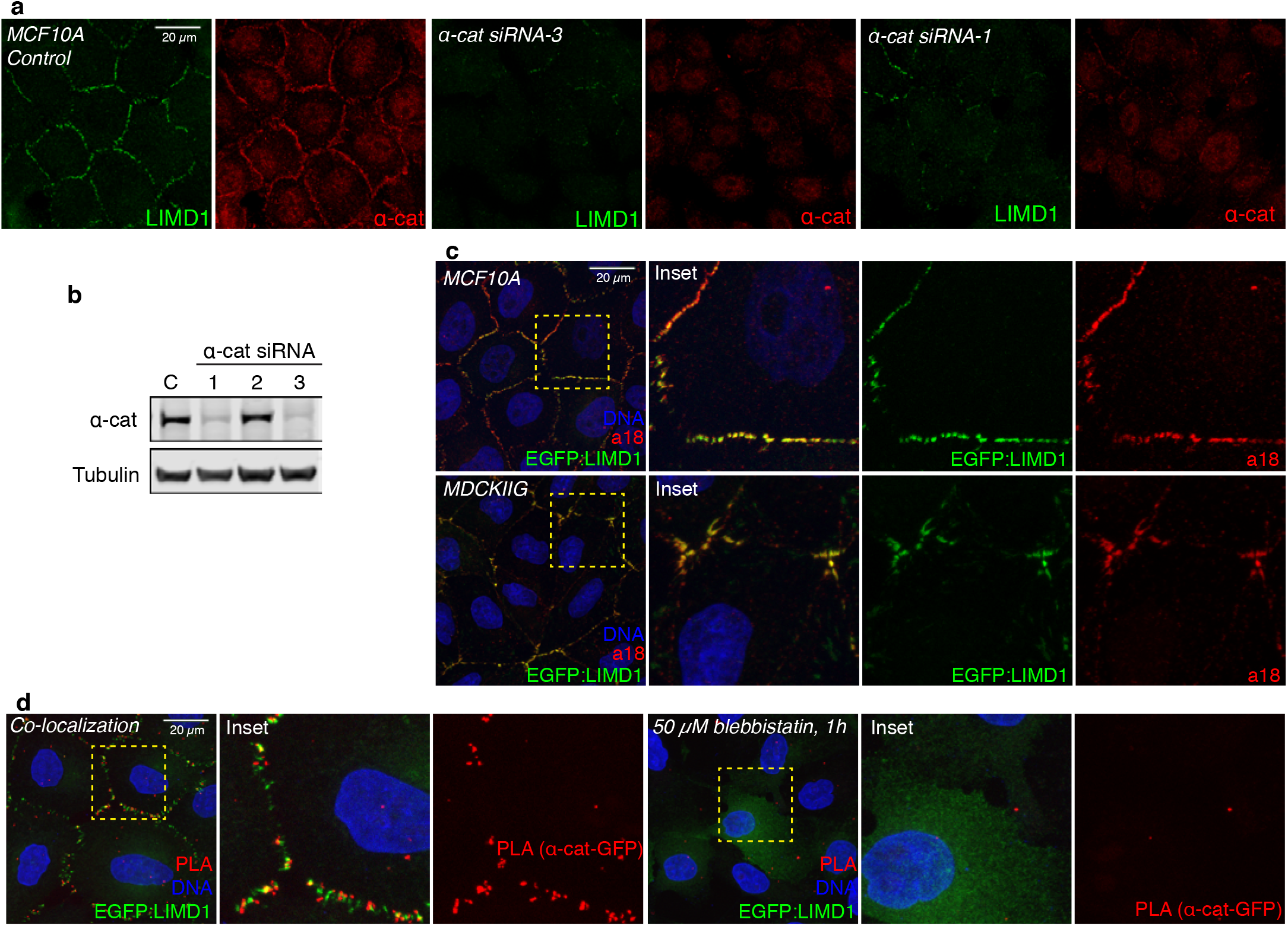
α-catenin is required for LIMD1 recruitment to adherens junctions. **(a)** MCF10A cells from the indicated cell lines were plated at low-density (15,000 cells/cm^2^) and cultured for 48 hours in total. Cells were transfected with control or α-catenin siRNAs, fixed in the presence of 0.5% Triton X-100 and then stained with LIMD1 (Millipore, green) and α-catenin (α-cat, red). **(b)** Western blot showing the knockdown efficiency of the α-catenin siRNAs tested. **(c)** MCF10A and MDCKIIG EGFP:LIMD1 cells grown at low density and stained with a18 (red) antibody and DNA (Hoechst, blue). Insets show higher magnification of the boxed regions. **(d)** Proximity ligation assay. MCF10A EGFP:LIMD1 cells were grown at low-density, treated with DMSO (control) or with 50 μM blebbistatin. Cells were induced with Dox for 24 h, fixed and incubated with mouse LIMD1 (Millipore) and mouse GFP antibodies detected with anti-mouse MINUS and anti-rabbit PLUS probes. Insets show higher magnification of the boxed regions.

### Tension-dependent localization of Lats proteins in mammalian epithelial cells

Published studies have described a variety of localizations for LATS1, including nuclear, cytoplasmic, and junctional (Jagannathan et al., 2016; Li et al., 2014; Szymaniak et al., 2015; Yin et al., 2013), and localization of LATS2 has not been reported. Whether their localization is affected by cytoskeletal tension has also not been described. To define their localization in mammalian epithelial cells, we created constructs for inducible expression of EGFP-tagged human LATS1 and LATS2 in lentiviral vectors. MCF10A and MDCKIIG cell lines with relatively homogeneous expression of EGFP:LATS1 were selected, but cells transduced with EGFP:LATS2 constructs tend to be lost during culture, even without addition of Dox, and we were only able to achieve patchy and transient expression of EGFP:LATS2 in both MDCKIIG and MCF10A cell lines. Moreover, we found that much higher levels of Dox were needed to induce visible EGFP:LATS2 expression than for any of our other cell lines (Table 1); these observations suggest that there is a strong selection against LATS2 expression, likely due to inhibition of YAP/TAZ activity. We also assayed several commercially available anti-Lats sera and confirmed that one of them has high specificity for Lats proteins. Western blotting of cells transduced to express EGFP:LATS1 or EGFP:LATS2 revealed that this antibody specifically recognizes human LATS1 (Fig. S4c). Moreover, the single band identified by this antibody in MCF10A cells was substantially reduced by LATS1 siRNA knockdown (Table 2 and Fig. S4d). Immunostaining of MCF10A cells with this antibody revealed a localization pattern similar to that of EGFP:LATS1, and this pattern is lost when cells were transfected with LATS1 siRNAs, but not with LATS2 siRNAs (Fig. S4f). By using canine-specific siRNAs, we found that this antibody appears to recognize both endogenous Lats1 and Lats2 on western blots of MDCKIIG cell lysates (Fig. S4e), and reveals a localization pattern in MDCKIIG cells similar to that of GFP-tagged human Lats proteins (Figs 3,4). By using LATS2 siRNA knockdown (Table 2), we also found an antibody that specifically recognized endogenous human LATS2 in Western blots, although not in immunostainings (Fig. S4d,g). Western blotting lysates of stably transduced cells identified conditions under which induced expression of EGFP:LATS1 was comparable to endogenous LATS1 expression (Fig. S4h-i), and these conditions were used for analysis of Lats protein localization. Direct comparisons to endogenous LATS2 levels were not possible due to the small fraction of EGFP:LATS2 expressing cells.

**Figure 3.**
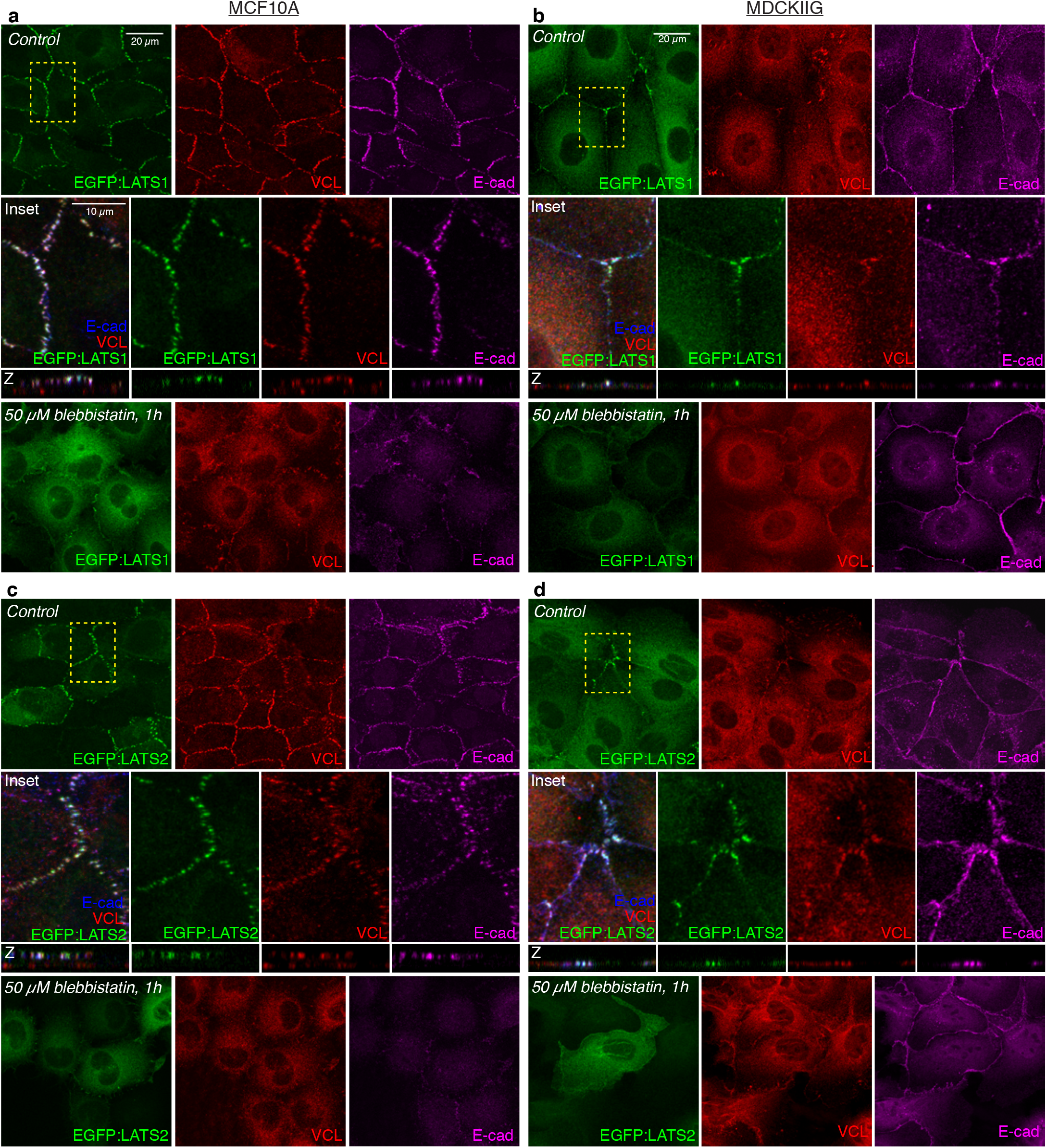
Tension-dependent localization of LATS proteins in MDCKIIG and MCF10A cell lines. **(a, c)** MCF10A or **(b, d)** MDCKIIG cells from the indicated cell lines were plated at low-density (15,000 cells/cm^2^) and cultured for 48 hours in total. Transgene expression was induced by adding Dox (Table 1) for 24 hours before treatment. Cells were treated with DMSO (Control) or 50 μM blebbistatin for 1 hour and then fixed in the presence of 0.5% Triton X-100 (**a, b**: MCF10A) or without detergent (**c, d**: MDCKIIG) and stained for VCL (red) and E-cad (blue/magenta). Square images are Z-projections (MDCKIIG) or apical slices (MCF10A) and are representative of at least 3 biological replicates. Insets show higher magnification of the boxed regions (yellow dashes). Vertical slices (Z) are shown as the same magnification as the boxed regions.

**Figure 4.**
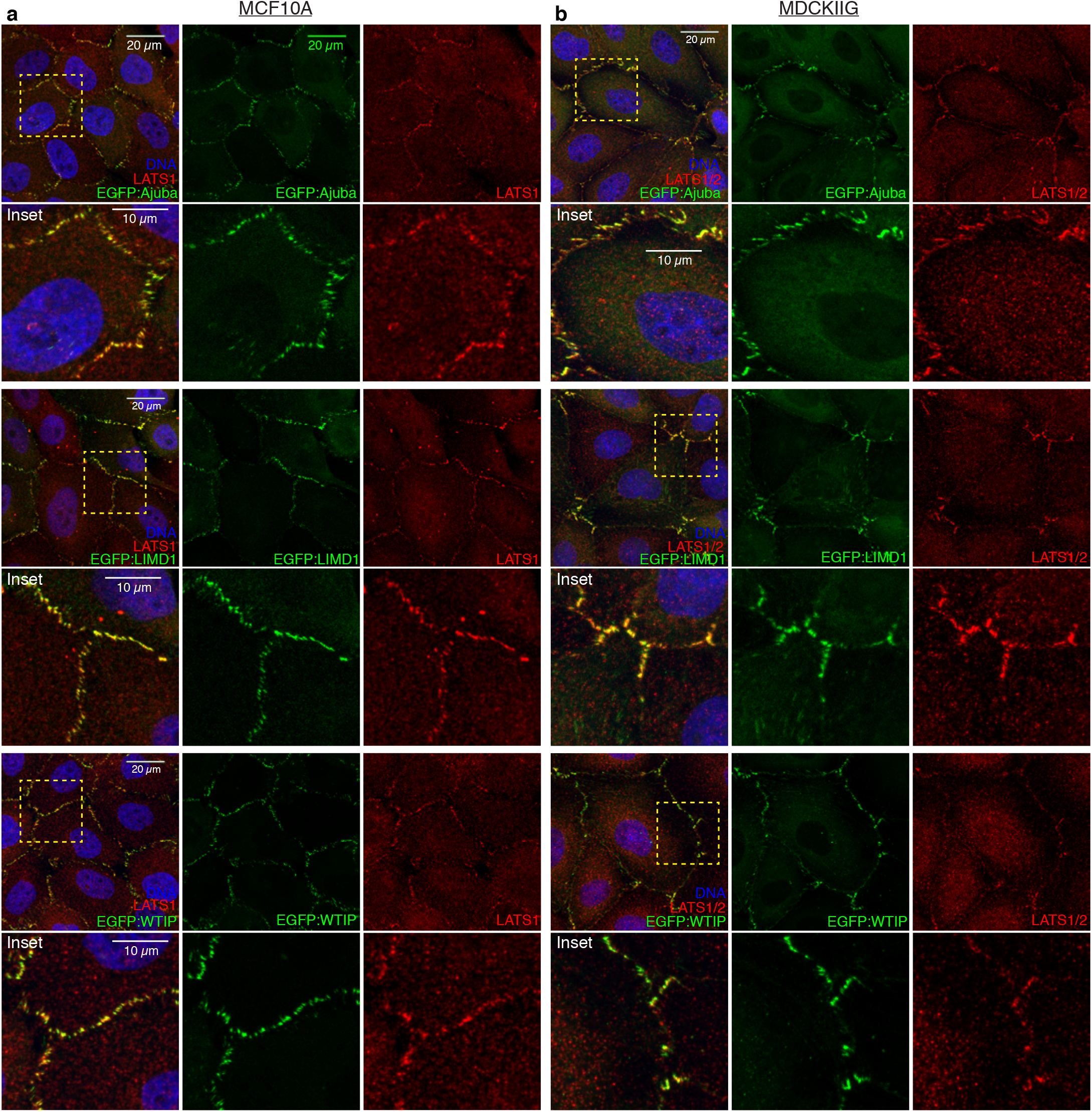
EGFP:Ajuba family proteins co-localize with LATS1. **(a)** MCF10A cells from the indicated stable cell lines were plated at low-density (15,000 cells/cm^2^) and cultured for 48 h in total. Transgene expression was induced with Dox (Table 1) for 24 h before fixation and then cells were stained for LATS1 (red) and DNA (Hoechst, blue). **(b)** MDCKIIG cells from the indicated stable cell lines were plated at low-density and treated as in (a). Insets show higher magnification of the boxed regions (yellow dashes).

**Table 2.**
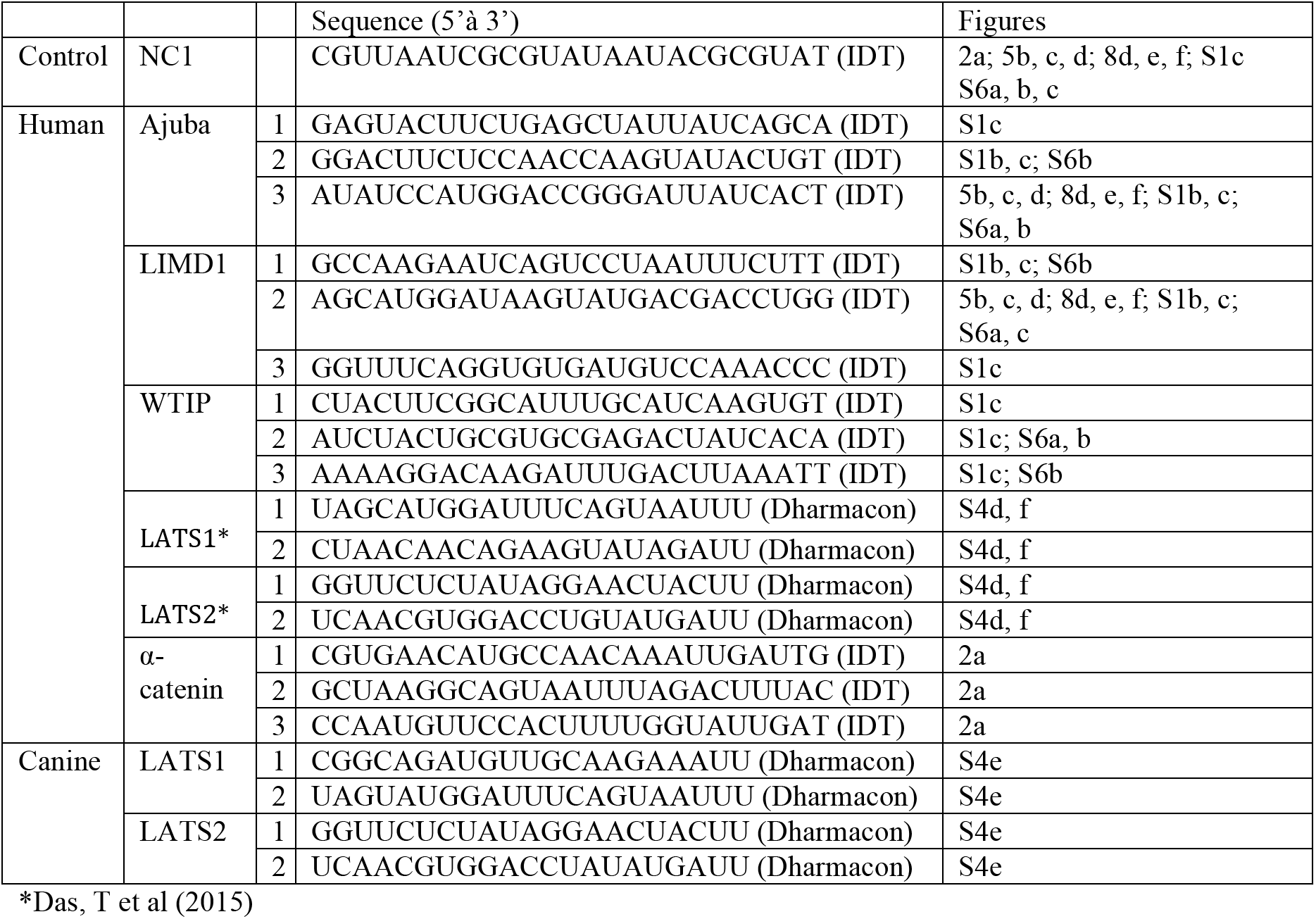
List of siRNAs used in this study.

LATS1 and LATS2 both accumulate in puncta overlapping adherens junctions in the same manner as the Ajuba family proteins (Figs 3, S4a-b). That is, in MDCKIIG cells they accumulate in puncta near intercellular vertices, whereas in MCF10A cells they accumulate in puncta around the circumference of the cell. However, in contrast to the Ajuba family proteins, accumulation of LATS proteins was never detected at focal adhesions (Fig. S5a-d). LATS puncta at adherens junctions overlap puncta of junctional VCL accumulation, implying that their localization is tension-dependent (Fig. 3). This was confirmed by observations that treatment of cells with either blebbistatin or Y-27632 leads to loss of the bright puncta of junctional Lats proteins, both in MCF10A and in MDCKIIG cells (Figs. 3, S5). Under these low-tension conditions, LATS proteins accumulate in the cytoplasm, but as for Ajuba family proteins we could also detect some low level, uniform localization around the cell circumference in MDCKIIG cells (Figs. 3b,d, S5). Thus, junctional localization of both LATS proteins in mammalian epithelial cells is modulated by cytoskeletal tension.

### LIMD1 recruits LATS to adherens junctions

The similar localization of Ajuba and Lats family proteins to puncta overlapping VCL at adherens junctions implies that they co-localize. To examine this directly, we used anti-LATS1 sera to stain cells expressing GFP-tagged Ajuba family proteins. This revealed extensive colocalization of LATS1 with each of the GFP-tagged Ajuba family proteins in puncta at cell junctions, both in MDCKIIG and MCF10A cells (Fig. 4). To further analyze this co-localization, we used PLA to probe the physical proximity of Ajuba and Lats family proteins. Using an anti-LIMD1 antibody in combination with an anti-GFP antibody on EGFP:LATS1-expressing cells, a clear and consistent PLA signal was detected in puncta along cell-cell junctions (Fig. 5a), which indicates that LIMD1 and LATS1 are closely associated on cell-cell junctions, most likely in direct contact. This PLA signal is lost upon blebbistatin treatment (Fig. 5a). No determination could be made regarding AJUBA or WTIP proximity to LATS1 because these antisera were not of sufficiently high quality to give a clear PLA signal.

**Figure 5.**
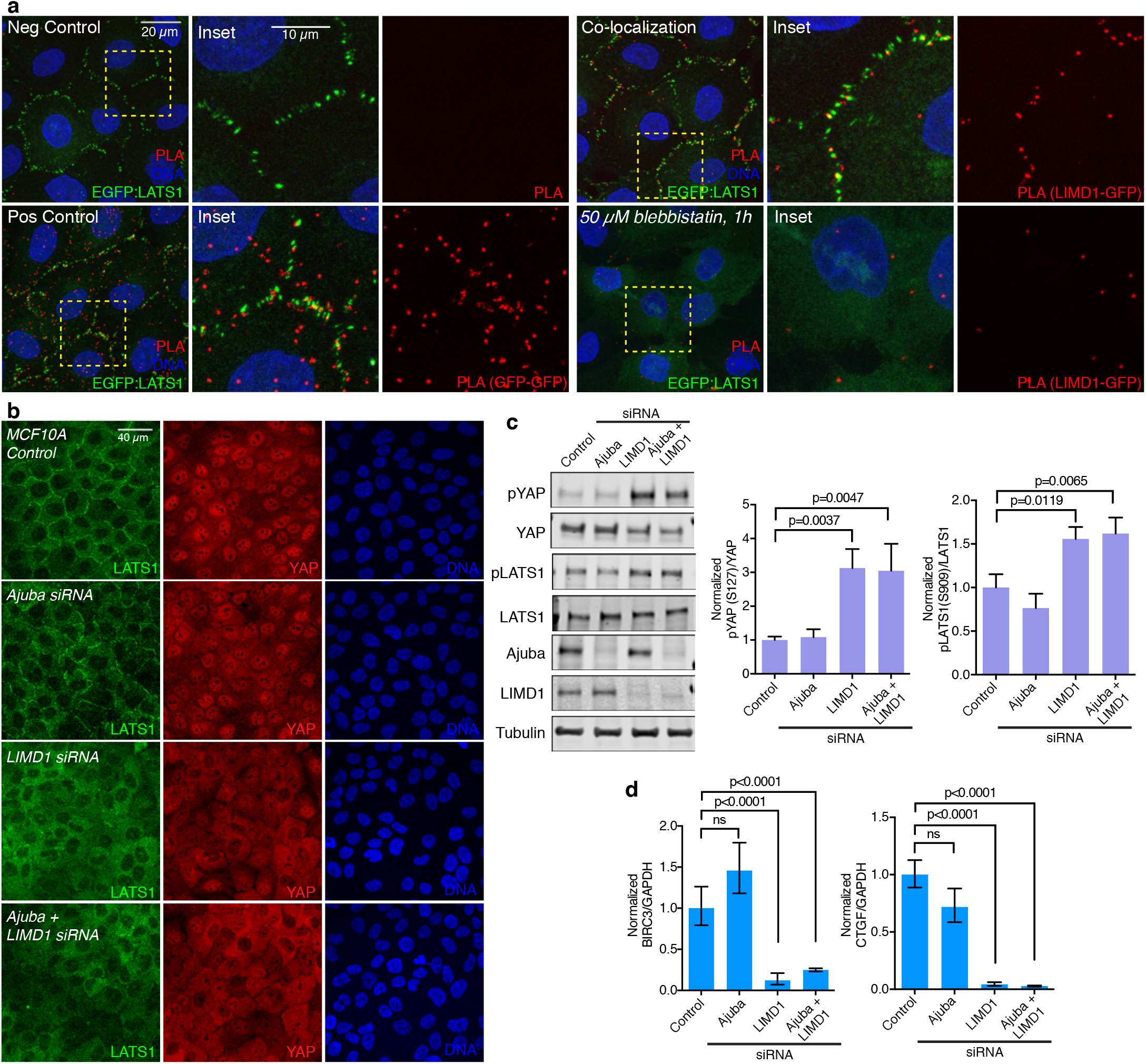
LIMD1 recruits LATS1 to adherens junctions and promotes YAP activity. **(a)** Proximity ligation assay. MCF10A EGFP:LATS1 cells were grown at low-density, induced with Dox and treated with or without 50μM blebbistatin. Cells were fixed and incubated with only secondary antibodies (*negative control*), mouse GFP antibody and both anti-mouse MINUS and PLUS probes *(positive control)*, and rabbit LIMD1 and mouse GFP antibodies detected with anti-mouse MINUS and anti-rabbit PLUS probes (*co-localization* and *50μM blebbistatin, 1h*). Insets show higher magnification of the boxed regions (yellow dashes). **(b)** MCF10A cells grown at low density transfected with control, Ajuba, LIMD1 or Ajuba+LIMD1 siRNAs, and stained for LATS1 (green), YAP (red) and DNA (Hoechst, blue). Images are Z-projections and representative of at least 3 biological replicates. **(c)** Western blots on lysates of MCF10A cells transfected with control, AJUBA, LIMD1 and Ajuba+LIMD1 siRNA, blotted using the indicated antisera. All the blots are from the same experiment and a representative loading control (Tubulin) is shown. Histograms show quantitation of the pYAP (S127) over YAP ratio and pLATS1 (S909) over LATS1 ratio from 3 biological replicates normalized to the ratio in the control siRNA transfection. Error bars indicate SEM and p-values less than 0.05 are shown. **(d)** Quantification of BIRC3 and CTGF mRNA abundance (n = 3 biological replicates) by RT-PCR on MCF10A cells transfected with control, Ajuba, LIMD1 and Ajuba+LIMD1 siRNA. Glyceraldehyde-3-phosphate dehydrogenase (GAPDH) was used as a standard. The BIRC3 or CTGF over GAPDH ratio was normalized to the ratio in the control siRNA transfection. Error bars indicate CI and p-values less than 0.05 are shown.

To determine whether the co-localization of Ajuba and Lats family proteins reflects a functional requirement for Ajuba proteins in recruiting Lats proteins to junctions, we examined LATS1 and LATS2 localization in MCF10A cells treated with at least two independent siRNAs targeting AJUBA, LIMD1, and/or WTIP (Table 2). Western blotting and immunostaining confirmed that these siRNAs could effectively reduce Ajuba family protein levels, and also could reduce the levels of the EGFP-tagged Ajuba family proteins (Fig. S1b,c, S6a). Knockdown of AJUBA or WTIP had no visible effect on LATS1 localization (Figs. 5b, S6b). In contrast, knockdown of LIMD1 clearly suppressed LATS1 and LATS2 recruitment to junctions (Figs 5b, S6b-c). Thus, junctional localization of Lats proteins in MCF10A cells specifically requires LIMD1.

When over-expressed, each of the three Ajuba family proteins has been reported to be able to co-immunoprecipitate with Lats proteins (Das Thakur et al., 2010; Sun and Irvine, 2013). Our observation that localization of endogenous LATS to junctional puncta in MCF10A cells specifically requires LIMD1 provided an opportunity to investigate the relationship between LATS protein localization and YAP activity. YAP activity was examined by assaying YAP localization, YAP phosphorylation, and expression of YAP target genes. Based on all three criteria, we found that YAP activity was reduced in MCF10A cells by knock-down of LIMD1, as LIMD1 siRNA reduced nuclear localization of YAP, increased phosphorylation of YAP at the key LATS phosphorylation site (S127), and reduced mRNA levels of YAP targets CTGF and BIRC3 (Figs 5b-d, S6b). Conversely, knockdown of AJUBA or WTIP had no effect on YAP activity (Figs 5b-d, S6b). We also assayed for potential additive effects of knockdowns of two Ajuba family genes, in all three pairwise combinations, but no additive effects were detected (Figs 5b-d, S6b). Staining western blots with anti-phospho-LATS sera revealed that knockdown of LIMD1, but not of AJUBA, increases LATS activation (Fig. 5c), which could account for the influence of LIMD1 on YAP activity. Altogether, these observations imply that amongst the three Ajuba family proteins, LIMD1 specifically regulates YAP activity in MCF10A cells, and that it does so by recruiting LATS to adherens junctions to suppress their activation.

### Regulation of Ajuba and Lats family proteins by cell density

Hippo signaling is influenced by cell density, and plays a key role in the density-dependent regulation of cell proliferation (contact-inhibition) (Zhao et al., 2007). To investigate the potential for junctional localization of Ajuba and Lats family proteins to contribute to density-dependent regulation of Hippo signaling, we investigated whether their localization is affected by cell density. At low densities (15,000 cells/cm^2^, corresponding to the conditions used in all of the experiments described above), Ajuba and Lats family proteins localize in bright puncta overlapping adherens junctions (Figs 1–4). In contrast, at high densities (150,000 cells/cm^2^) bright puncta of junctional localization are rare or absent (Figs. 6, S7).

Examination of YAP activity in MCF10A cells confirmed that changes in YAP correlate with these changes in LIMD1 and LATS protein localization. Thus, under our low cell density conditions, YAP is predominantly nuclear, whereas under our high cell density conditions, YAP is predominantly cytoplasmic (Figs 6, S7). (We note that expression of EGFP:LATS2 at detectable levels reduced YAP nuclear localization at low cell density, but further reductions are nonetheless evident at higher cell density, Fig. 6). The correlation between YAP localization and LIMD1 and LATS localization suggests that density-dependent regulation of their localization could contribute to density-dependent regulation of YAP. Consistent with this hypothesis, we observed that LIMD1 knockdown decreases YAP activity at low cell density, but has no significant effect at high cell density (Figs 5b-d, 8e-f).

**Figure 6.**
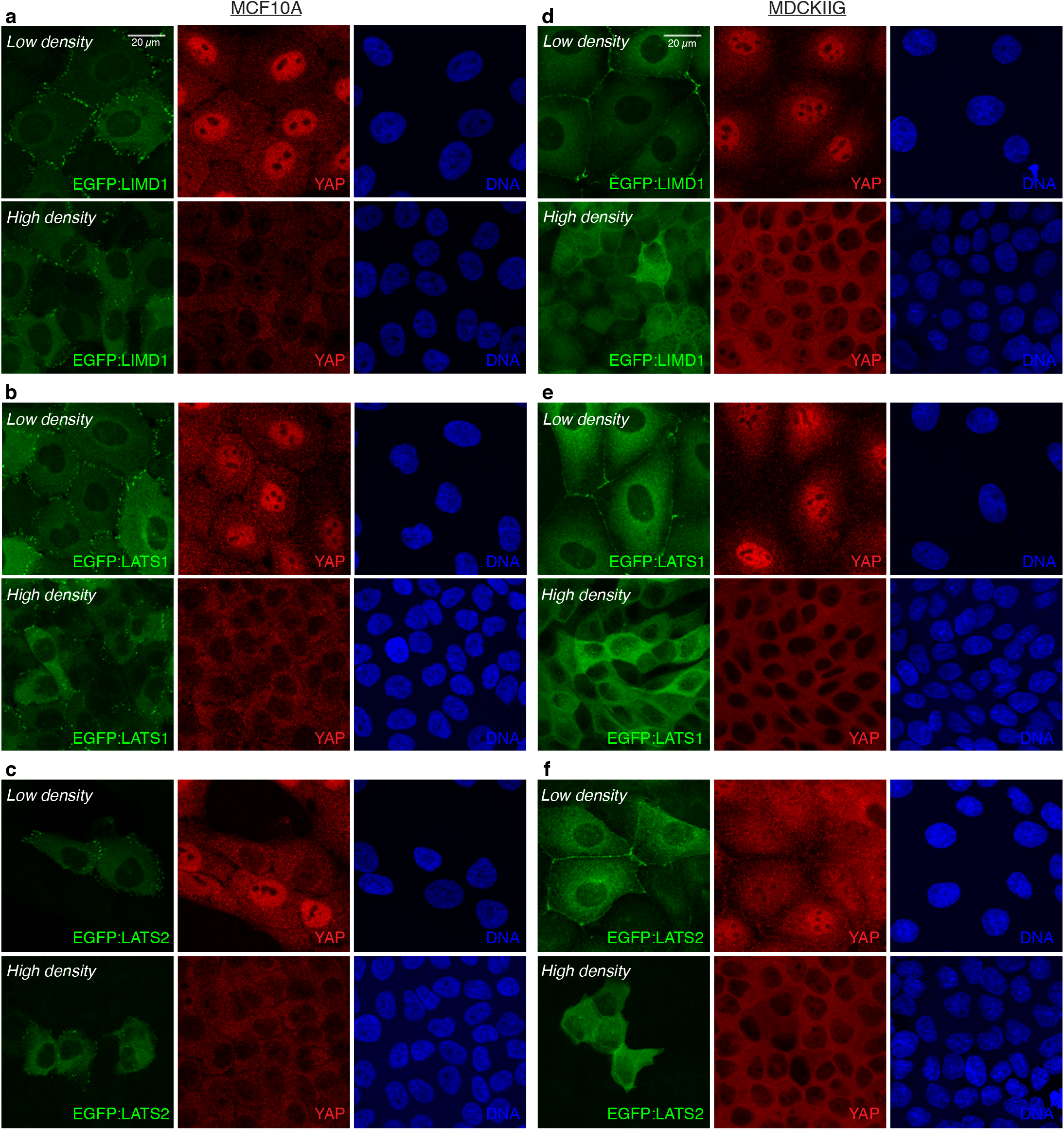
Regulation of LIMD1 and LATS proteins localization and YAP activity by cell density. **(a, b, c)** MCF10A or **(d, e, f)** MDCKIIG from the indicated cell lines were plated at low (15,000 cells/cm^2^) and high density (150,000 cells/cm^2^) and grown for 48 hours in total. Transgene expression was induced by adding Dox 24 hours before fixation (Table 1). Cells were stained with YAP (red) and DNA (Hoechst, blue). Images are projections through Z stacks.

The localization of Ajuba and Lats family proteins at high density becomes similar to that observed when myosin activity is inhibited at low cell density: junctional localization is reduced in MCF10A cells, and a weak, uniform junctional localization is observed in MDCKIIG cells (Figs. 6, S7). Indeed, examination of phospho-myosin (pMLC) and VCL localization indicate that cytoskeletal tension is lower in cells at high cell density (Fig. S7c-f), which could account for the observed changes in Ajuba and Lats family protein localization. Moreover, the tension-dependence of Ajuba and Lats family protein localization also correlates with influences of cytoskeletal tension on YAP activity. Thus, under the same doses of blebbistatin that Ajuba and Lats family proteins are lost from puncta at adherens junctions despite low cell density, LATS phosphorylation is increased, and YAP activity is reduced, as revealed by Western blotting with anti-phospho-YAP and immunostaining for YAP (Fig. 7).

**Figure 7.**
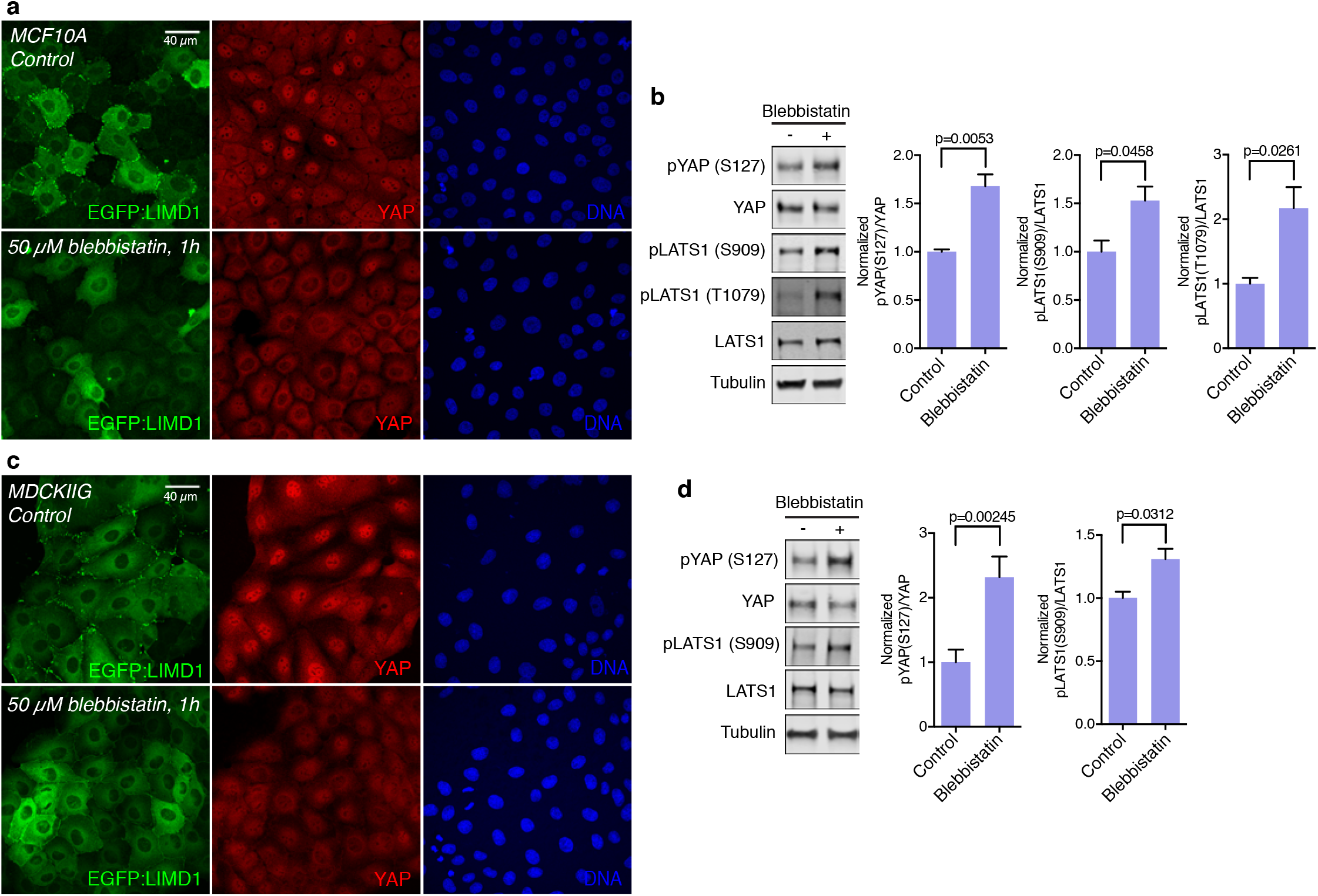
Tension-dependence of Ajuba and Lats family protein localization correlates with influences of cytoskeletal tension on YAP activity. **(a)** MCF10A EGFP:LIMD1 cells were plated at low density and grown for 48 hours in total. Transgene expression was induced by adding Dox 24 h before treatments (Table 1). Cells were treated with DMSO (control) or treated with 50 μM blebbistatin for 1 hour. After the treatment, cells were stained for YAP (red) and DNA (Hoechst, blue) **(b)** Western blots on lysates of MCF10A cells treated with DMSO (control) or 50 μM blebbistatin for 1 hour, blotted using the indicated antisera. All the blots are from the same experiment and a representative loading control (Tubulin) is shown. Histograms show quantitation of the pYAP (S127) over YAP ratio and pLATS1 (S909 and T1079) over LATS1 ratio from 3 biological replicates normalized to the ratio in the control. **(c)** MDCKIIG EGFP:LIMD1 cells were grown and stained as in **(a)**. Images are Z-projections and representative of at least 3 biological replicates. **(d)** Western blots on lysates of MDCKIIG cells treated with DMSO (control) or 50 μM blebbistatin for 1 hour, blotted using the indicated antisera. Tubulin was used as a loading control. Histograms show quantitation of the pYAP (S127) over YAP ratio and pLATS1 (S909) over LATS1 ratio from 3 biological replicates normalized by to the ratio in the control. In all histograms, error bars indicate SEM and p-values less than 0.05 are shown.

### Ajuba and Lats family proteins contribute to Rho-mediated regulation of Hippo signaling

Multiple upstream inputs into Hippo signaling appear to act through, or are dependent upon, the small GTPase Rho. For example, YAP/TAZ activation associated with cell attachment to substrates, large cell area, stiff substrates, or activation of certain G protein-coupled receptor (GPCR) signaling pathways is suppressed by Rho inhibitors (Dupont et al., 2011; Yu et al., 2012; Zhao et al., 2012), whereas expression of activated-Rho can promote YAP activation (Zhao et al., 2012). However, the mechanism(s) by which Rho activity ultimately impinges upon YAP activity have remained unclear. Based upon the well-established ability of Rho to influence F-actin and myosin activity (Fig. S8e) (Etienne-Manneville and Hall, 2002), we asked whether Rho-mediated regulation of Hippo signaling involves Ajuba family proteins.

This was examined using MCF10A cells plated at high density, which have low recruitment of Ajuba and Lats family proteins to adherens junctions, and correspondingly low YAP activity (Figs. 8, S8). Endogenous Rho proteins were activated either by treating cells with lysophosphatidic acid (LPA), which activates Rho through GPCRs (Yu et al., 2012), or Rho activator II (cytotoxic necrotizing factor-1 (CNF1) from *E. coli*), which deamidates glutamine-63 of RHOA and its homologs, preventing GTP hydrolysis and thereby keeping Rho proteins in their GTP-bound, active state (Schmidt et al., 1997). Each of these treatments substantially decreased LATS activity, and increased YAP activity (Figs 8, S8). At the same time, Rho activation substantially increased localization of Ajuba and Lats family proteins to adherens junctions (Figs 8, S8). Thus, Rho-mediated YAP activation correlates with Rho-mediated Ajuba and Lats protein localization to cell-cell junctions.

**Figure 8.**
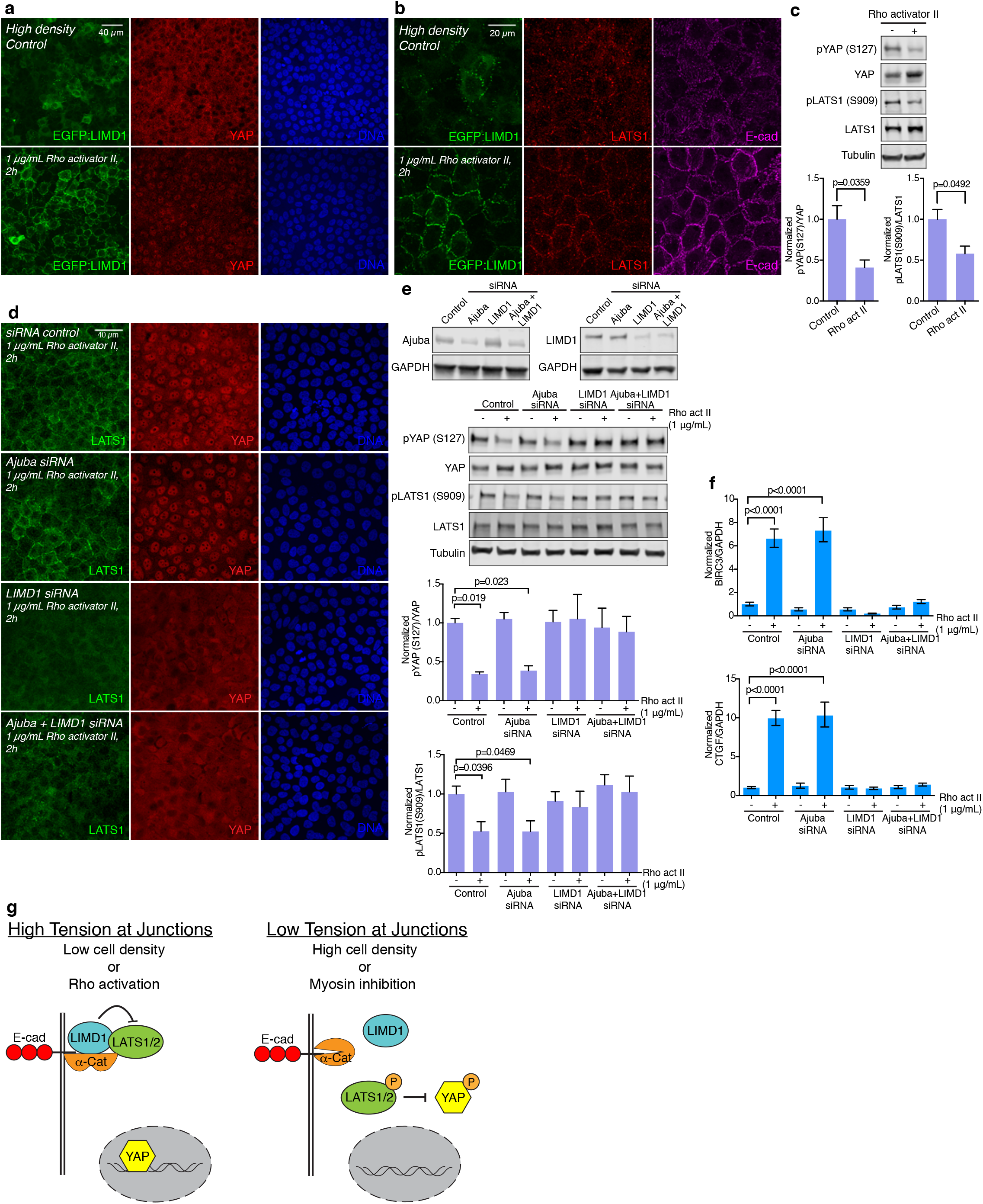
Ajuba and Lats family proteins contribute to Rho-mediated regulation of Hippo signaling. **(a)** MCF10A EGFP:LIMD1 cells grown at high density, induced with Dox for 24 h before treatments (Table 1) and mock-treated with water (control) or treated with 1 μg/mL Rho activator II for 2 hours. After treatment, cells were stained for YAP (red) and DNA (Hoechst, blue) **(b)** Same cell line and treatments as in **(a)**, but cells were stained for LATS1 (red) and E-cad (magenta). Images are Z-projections and representative of at least 3 biological replicates. **(c)** Western blots on lysates of MCF10A cells mock-treated with water (control) or treated with 1 μg/mL Rho activator II for 2 hours, blotted using the indicated antisera. All the blots are from the same experiment and a representative loading control (Tubulin) is shown. Histograms show quantitation of the pYAP (S127) over YAP ratio and pLATS1 (S909) over LATS1 ratio from 3 biological replicates normalized to the ratio in the control. Error bars indicate SEM and p-values less than 0.05 are shown. **(d)** MCF10A cells grown at high density and transfected with control, Ajuba, LIMD1 or Ajuba+LIMD1 siRNA. After 48 hours, cells were treated with 1 μg/mL Rho activator II for 2 hours and stained for LATS1 (green), YAP (red) and DNA (Hoechst, blue). Images are Z-projections and representative of at least 3 experiments. **(e)** Western blots on lysates of MCF10A cells grown at high density transfected with control, Ajuba, LIMD1 or Ajuba+LIMD1 siRNA, blotted using the indicated antisera. Efficiency of the knockdown is reduced when the cells are transfected at high density as compared to low density (compare with Figure 3 supplement 1). All the blots are from the same experiment and a representative loading control (Tubulin or GAPDH) is shown. Histograms show quantitation of the pYAP (S127) over YAP ratio and pLATS1 (S909) over LATS1 ratio from 3 biological replicates normalized to the ratio in the control siRNA transfection. Error bars indicate SEM and p-values less than 0.05 are shown. **(f)** Quantification of BIRC3 and CTGF mRNA abundance (n = 3 biological replicates) by RT-PCR on MCF10A cells transfected with control, Ajuba, LIMD1 or Ajuba+LIMD1 siRNA and treated with 1 μg/mL Rho activator II. Glyceraldehyde-3-phosphate dehydrogenase (GAPDH) was used as a standard. The BIRC3 or CTGF over GAPDH ratio was normalized to the ratio in the control siRNA transfection. Error bars indicate CI and p-values less than 0.05 are shown. **(g)** Summary model for regulation of YAP by LIMD1. When the cells are under high tension (left, e.g. cell low density or Rho activation), LIMD1 is associated with adherens junctions through α-catenin, where it recruits and inhibits LATS. This allows YAP to go to the nucleus and activate transcription. When cells are under low tension (right, e.g. high density or myosin inhibition), the α-catenin conformation is altered, LIMD1 and LATS are released from junctions, and LATS can be activated. Activated (phosphorylated) LATS phosphorylates and inhibits YAP by promoting its cytoplasmic localization and degradation.

To directly test the requirement for Ajuba family proteins, and the significance of the correlation between LATS recruitment to junctions and activation of YAP in response to Rho activation, we examined the consequences of siRNA-mediated knockdown AJUBA and/or LIMD1 on cells plated at high density and then treated with Rho activator II. These experiments revealed that knock down of AJUBA did not visibly decrease Rho-mediated LATS1 recruitment or YAP activation (Fig. 8). Conversely, knockdown of LIMD1 suppressed Rho-mediated LATS1 recruitment to junctions, YAP activation and expression of YAP target genes (Fig. 8). Thus, LIMD1 is specifically required for Rho-mediated LATS re-localization and YAP activation.

## DISCUSSION

Hippo signaling has emerged as a key conduit for transduction of cells’ responses to their biomechanical environment. Nonetheless, the molecular mechanisms by which biomechanical signals are perceived and transduced to influence YAP and TAZ activity in mammalian cells have remained poorly understood. Our observations that LIMD1 localizes to puncta at adherens junctions under conditions of low cell density, Rho activation, and cytoskeletal tension, that this LIMD1 recruits Lats kinases to adherens junctions and inhibits Lats activity, and that LIMD1 is required for YAP activation under conditions of low cell density, Rho activation, and cytoskeletal tension, all support a model in which tension-dependent recruitment and inhibition of Lats kinases in complexes with LIMD1 at adherens junctions contributes to biomechanical (and biochemical) regulation of Hippo signaling (Fig. 8g).

The mechanism by which LIMD1 inhibits LATS activation is not clear, but studies of Jub inhibition of Warts in *Drosophila* suggested a model in which recruitment of Warts to Jub sequesters it away from upstream complexes that promote Warts activation (Sun et al., 2015). In *Drosophila* these include junctional Expanded complexes, where Hippo, Salvador, and Expanded can be seen to overlap with activated-Warts (Sun et al., 2015), and apical Merlin complexes (Su et al., 2017). Junctional or apical localization of proteins that could scaffold assembly of Lats-activating complexes has also been reported in mammalian cells (Sun and Irvine, 2016), and observations that at least in some contexts the key upstream regulators Merlin and Angiomotins associate with tight junctions (Yi et al., 2011) suggests that sequestration of LATS into adherens junction complexes might prevent its activation. It is also possible that association with LIMD1 physically precludes association of LATS with one or more upstream activators.

The absence of visible recruitment of Lats proteins to focal adhesions, despite the presence of Ajuba family proteins indicates that there must be additional cues at adherens junctions that promote the recruitment of Lats kinases. The absence of Lats kinases at focal adhesions also implies that distinct mechanisms are involved in biomechanical signaling mediated through focal adhesions. This is consistent with observations that some biomechanical signals that impinge on focal adhesions, such as substrate stiffness, substrate stretching, or increasing cell shape through modulation of cell attachment sites, can influence YAP/TAZ activity through mechanisms that are independent of LATS activity (Aragona et al., 2013; Das et al., 2016; Dupont et al., 2011). Similarly, a reported lack of requirement for Ajuba family proteins for “mechanical signals” is not inconsistent with our results, because that study (Jagannathan et al., 2016) examined potential requirements for AJUBA and LIMD1 in YAP activation under conditions of stiff substrates or cell spreading, which are expected to exert tension on focal adhesions rather than adherens junctions. Jagannathan et al. (2016) also reported observing increased LATS activity only when both AJUBA and LIMD1 were knocked down, and not when LIMD1 alone was knocked down. However, this was under low density conditions differed from ours, as they examined isolated cells lacking cell-cell junctions, which consequently could not have the LIMD1-specific junctional recruitment of LATS that we identified. We also note that while Jagannathan et al. (2016) reported that LIMD1 and LATS1 could be detected at adherens junctions in cells cultured at “high cell density”, based on cell area some of their high density images appear to correspond conditions that we would classify as low cell density.

Our observations implicate contact inhibition of cell proliferation (more accurately described as cell density-dependent regulation of cell proliferation (Puliafito et al., 2012)) as a biological context where tension-dependent recruitment of LIMD1 and LATS to adherens junctions plays a role in modulating Hippo signaling. Hippo signaling has been shown to mediate density-dependent regulation of cell proliferation (Zhao et al., 2007), but the mechanism by which this occurs were unclear. The reduction in junctional localization and LATS inhibition as cell density increases appears to stem from decreased cytoskeletal tension, as it correlates with reduced phospho-myosin staining and VCL localization, and it can be reversed by activation of Rho. Conversely, pharmacological reduction of cytoskeletal tension can abolish junctional localization of LIMD1 and LATS even under low cell density conditions. The observation that cytoskeletal tension is reduced as cells become more crowded is also consistent both with modeling of epithelial mechanics (Noll et al., 2017; Pan et al., 2016; Puliafito et al., 2012), and with experimental studies in *Drosophila*, where growth-induced cell crowding has been shown to reduce cytoskeletal tension (Pan et al., 2016).

Our observations also imply that formation of junctional complexes between LATS and LIMD1 contributes to Hippo pathway regulation by GPCR signaling. Regulation of Hippo signaling by GPCRs is known to depend upon Rho, but the molecular mechanism by which Rho affects Hippo signaling has been unclear. GPCR signaling is activated by LPA, and mediated through Rho activation (Yu et al., 2012), both of which we found promote junctional recruitment of LIMD1 and LATS, and LIMD1-dependent activation of YAP.

Our studies identified all three Ajuba family proteins as exhibiting tension-dependent recruitment to adherens junctions. The distinct localization profiles observed in MCF10A versus MDCKIIG cells are consistent with tension-dependent localization, as the distinct organization of the cytoskeleton in cells with punctate versus linear adherens junctions is expected to correlate with differences in the spatial pattern of tension along junctions (Takeichi, 2014). This tension-dependent recruitment is likely mediated through conformational changes in α-catenin structure (Yonemura et al., 2010), as both AJUBA and Jub have been reported to associate with α-catenin (Marie et al., 2003; Rauskolb et al., 2014), and the puncta of Ajuba family localization at adherens junctions co-localize both with VCL, which associates with stretched α-catenin, and with a monoclonal antibody (a18) that specifically recognizes stretched α-catenin. Junctional localization of LIMD1 also requires α-catenin, and we confirmed that LIMD1 and α-catenin are closely associated at junctions using PLA. As Ajuba family proteins have many other functions besides their contribution to Hippo signaling (Schimizzi and Longmore, 2015), our observation that the localization of all three proteins is regulated by tension raise the possibility that some of their other functions could also be modulated by cytoskeletal tension.

Given their similar tension-dependent recruitment, and the ability of all three Ajuba family proteins to associate with Lats family proteins (Das Thakur et al., 2010), the specific requirement for LIMD1 in LATS recruitment was unexpected. Although there could be in vivo differences in requirements due to different expression patterns, both AJUBA and LIMD1 are clearly expressed in the MCF10A cells we examined. It’s conceivable that there that exist differences in binding affinity, differences in association with partner proteins, or differences in post-translational modifications, which account for the specific requirement for LIMD1 as opposed to other Ajuba family proteins, and this will be an important question for future studies.

## ACKNOWLEDGEMENTS

We thank W.J Nelson, J. Debnath, G. Sun, A. Nagafuchi, N. Yabuta and H. Nojima for reagents. This research was supported by NIH grant R01 GM121537 (KDI) and NJCCR fellowship DHFS16PPC035 (CI).

## MATERIALS AND METHODS

### Plasmids

To generate Dox-inducible lentiviral vector, coding sequence of TurboRFP in pTRIPZ (GE Healthcare) was replaced with the following GFP-fusion proteins. EGFP-tagged human AJUBA, LIMD1, LATS1 and WTIP were generated by PCR from pcDNA3.1-V5:His containing AJUBA, LIMD1, LATS1 or WTIP (Reddy and Irvine, 2013; Sun and Irvine, 2013) and cloned into pEGFP-C3. Human EGFP-LATS2 was a gift of N. Yabuta and H. Nojima, Osaka University (Yabuta et al., 2011).

### Generation of stable cell lines

Lentiviral Dox-inducible vectors were co-transfected with Trans-Lentiviral Packaging Mix into HEK-293T (Dharmacon) cells using calcium phosphate. After 16 h of transfection, the medium was replaced with UltraCULTURE medium (Lonza) supplemented with 1X Gluta-MAX (Life Technologies) and antibiotics. Starting the next day, the lentivirus-containing supernatants were collected every 24 h for 2 days and stored at 4°C. After spin-down at 2,000 rpm for 10 min to remove cell debris, supernatants were filtered using a 0.45-μm-pore size filter. About 30 ml of filtered supernatants were then transferred into ultracentrifuge tubes containing 5 ml of 20% sucrose in PBS cushion. Lentiviral particles were concentrated by centrifugation at 100,000 *g* for 2 h at 4°C and resuspended in 1× HBSS buffer. MCF-10A or MDCKIIG cells were incubated overnight with lentiviral particles and 8 μg/ml polybrene (Sigma) in media without serum and replaced with complete medium the next morning. Antibiotics selection was started 2 days after transduction with 2 μg/ml puromycin and maintained on selection medium for an additional 7 days. After selection, cells were isolated to establish clonal populations. Induction of the transgenes was done with Dox 24 h before fixation or harvesting.

### Cell culture, transfections and treatments

MDCKIIG (a gift from W.J Nelson, Stanford University) cells were cultured in low glucose-Dubecco’s modified Eagle’s medium (DMEM) (Life Technologies) supplemented with 10% FBS and antibiotic-antimycotic, and MCF10A (a gift from Jay Debnath, UCSF) were cultured in DMEM/F-12 (Life Technologies) supplemented with 5% horse serum, epidermal growth factor (20 μg/ml), insulin (10 μg/ml), cholera toxin (0.1 μg/ml), hydrocortisone (0.5 μg/ml), and antibiotic-antimycotic at 37°C and 5% CO_2_. For immunostainings, cells were grown on coverslips coated with 0.6 mg/mL of collagen for 15 min at room temperature and washed with PBS.

For the cytoskeletal inhibitor treatments, cells were grown at low density (15,000 cells/cm^2^) for 48 hours and blebbistatin (50 μM) (Sigma) or Y-27632 (10 μM for MCF10A and 20 μM for MDCKIIG cells) (Cytoskeleton) were applied to the cells for 1 hour. For cytoskeletal activators, cells were grown at high density (150,000 cells/cm^2^) for 48 hours and Rho activator II (1 μg/ml; Cytoskeleton) or lysophosphatidic acid (LPA) (1 μM; Sigma) was added for 2 or 1 hour, respectively.

For siRNA delivery, cells were transfected with Lipofectamine RNAi Max (Life Technologies) according with the manufacturer’s protocols, and fixed or harvested 48 hours after transfection.

### Immunostaining and imaging

Cells were fixed with 4% paraformaldehyde in PBS++ (phosphate-buffered saline supplemented with 100 mM MgCl_2_ and 50 mM CaCl_2_) for 10 min at room temperature. For the detection of VCL, α-catenin or a18, cells were fixed in 1% paraformaldehyde with 0.5% Triton X-100 in PBS++ for 3 min, rinsed and then fixed in 1% PFA in PBS++ for 10 min. Then cells were washed three times for 5 min each with 200 mM glycine containing PBS, followed by permeabilization with 0.5% Triton X-100 in PBS for 20 min.

After blocking with 5%bovine serum albumin (BSA) in PBS for 1 h, cells were incubated with primary antibody diluted in 5% BSA/PBS overnight at 4°C. After washing with PBS, cells were incubated with Alexa Fluor 488- (Life Technologies), Cy3- or Alexa Fluor 647–conjugated secondary antibodies (Jackson ImmunoResearch) for 2 h and washed 4 times with PBS. Cell nuclei were counterstained with Hoechst 33342 (1 μg/ml; Invitrogen) and mounted with mounting media (Dako). Antibodies used for immunostaining include mouse anti-Yap (1:100; Santa Cruz Biotechnology, sc-101199), rabbit anti-Yap (1:100, Cell Signaling Technology, #14074), rabbit anti-Lats1 (1:600; Cell Signaling Technology, #3477), rabbit anti-α-catenin (1:500; Sigma, C8114), mouse anti-vinculin (1:2,000; Sigma, V9131), rat anti-a18 (1:50; gift from A. Nagafuchi), mouse anti-ZO-1 (1:1,000; Life Technologies, #33-9100), mouse anti-phospho-myosin light chain (S19) (1:200; Cell Signaling Technology, #3675), rat anti-E-cadherin (1:500; Life Technologies, #13-1900), rabbit anti-Ajuba (1:500; Cell Signaling Technologies, #4897), rabbit anti-LIMD1 (1:500; Bethyl, A303-182A), mouse anti-LIMD1 (1:500; EMD Millipore, MABD85). F-actin was stained with Alexa Fluor 647 phalloidin (1:50; Life Technologies). Images were acquired using LAS X software on a on a Leica TCS SP8 confocal microscope system using a HC PL APO 63x/1.40 objective. Images were processed using ImageJ and Adobe Photoshop CS6, and figures were assembled in Adobe Illustrator CS6.

### Proximity ligation assay

Proximity ligation assay (PLA) was done using Duolink^®^ Proximity Ligation Assay kit and performed according to manufacturer’s instructions (Sigma). Fixation and permeabilization steps were done as a normal immunostaining procedure. Antibodies used for PLA include mouse GFP (1:250; Life Technologies, A11120), rabbit LIMD1 (1:250; Bethyl, A303-182A), rabbit α-catenin (1:250; Sigma, C8114) and secondary anti-mouse MINUS and anti-rabbit PLUS probes were used. As a negative control of the PLA signal, only secondary antibodies were used and as a positive control, EGFP:LATS1 was recognized by using only GFP antibody and secondary anti-mouse MINUS and anti-mouse PLUS probes.

### Immunoblotting

Cells were lysed in lysis buffer [50 mM Tris-HCl (pH 7.4), 150 mM NaCl, 1%Triton X-100, 0.1% CHAPS, 0.1% NP-40, 1 mM EDTA, 5% glycerol] or in 2X Laemmli Sample Buffer (Bio-Rad) supplemented with protease inhibitor cocktail (Roche) and phosphatase inhibitor cocktail (Calbiochem). Protein samples were loaded in 4 to 15% gradient gels (Bio-Rad). Antibodies used for immunoblotting include rabbit anti-Lats1 (1:1000; Cell Signaling Technology, #3477), rabbit anti-Lats2 (1:1000; Bethyl, A300-479A), rabbit anti-phospho-Lats1 (T1079) (1:1000; Cell Signaling Technology, #8654), rabbit anti-phospho-Lats1 (S909) (1:1000; Cell Signaling Technology, #9157), rabbit anti-phospho-Yap (S127) (1:1000; Cell Signaling Technology, #4911), rabbit anti-Yap (1:1000; Abcam, ab52771), mouse anti-LIMD1 (1:1000; EMD Millipore, MABD85), rabbit anti-Ajuba (1:1000; Cell Signaling Technology, #4897), goat anti-WTIP (1:1000; Santa Cruz Biotechnology, sc-241738), rabbit α-catenin (1:1000; Sigma, C8114), mouse anti-GFP (1:1000; Cell Signaling Technology, #2955), rabbit anti-GFP (1:1000; Cell Signaling Technology, #2555). As loading controls, mouse anti-α-tubulin (1:10,000; Sigma, T6199), rabbit anti-GAPDH (1:5,000; Santa Cruz Biotechnology, sc-25778) or mouse anti-GAPDH (1:10,000; Novus Biologicals, NBP2-27103) were used. Blots were visualized and quantified with fluorescent-conjugated secondary antibodies (LI-COR Biosciences) and Odyssey Imaging System (LI-COR Biosciences). All the blots from the same experiment were loaded equally and run on multiple gels in parallel, then blotted with the indicated antibodies and loading controls to be able to make comparisons between phosphorylated protein and total protein.

### Quantitative PCR Protocol

Total RNA from MCF10A cells was extracted using Trizol reagent (Life Technologies) according to manufacturer’s instructions. Reverse transcription PCR (RT-PCR) was carried out on 2 μg of total RNA using High Capacity cDNA Reverse Transcription kit (Applied Biosystems) according to the manufacturer’s instructions. First strand cDNA was subjected to qPCR using SYBR Select Master Mix and specific BIRC3, CTGF and GAPDH (as housekeeping gene) primers.

### Statistical analysis

Statistical significance was determined using Graphpad Prism software, with a paired two-tailed *t* test for two sample comparisons or analysis of variance (ANOVA) for more than two samples, with p<0.05 set as the criteria for significance. The Tukey test was used to derive adjusted p-values for multiple comparisons. For image quantitation, statistical tests of differences were calculated on the log of the ratios. Error bars on figure panels show SEM, except for the ones from RT-PCR which show the 95% confidence interval (CI).

